# SARS-CoV-2 spike protein induces TLR-4-mediated long-term cognitive dysfunction recapitulating post-COVID syndrome

**DOI:** 10.1101/2022.06.07.495149

**Authors:** Fabricia L. Fontes-Dantas, Gabriel G. Fernandes, Elisa G. Gutman, Emanuelle V. De Lima, Leticia S. Antonio, Mariana B. Hammerle, Hannah P. Mota-Araujo, Lilian C. Colodeti, Suzana M. B. Araújo, Talita N. da Silva, Larissa A. Duarte, Andreza L. Salvio, Karina L. Pires, Luciane A. A. Leon, Claudia Cristina F. Vasconcelos, Luciana Romão, Luiz Eduardo B. Savio, Jerson L. Silva, Robson da Costa, Julia R. Clarke, Andrea T. Da Poian, Soniza V. Alves-Leon, Giselle F. Passos, Claudia P. Figueiredo

## Abstract

Cognitive dysfunction is often reported in post-COVID patients, but its underlying mechanisms remain unknown. While some evidence indicate that SARS-CoV-2 can reach and directly impact the brain, others suggest viral neuroinvasion as a rare event. Independently of brain viral infection, the ability of SARS-CoV-2 spike (S) protein to cross the BBB and reach memory-related brain regions has already been shown. Here, we demonstrate that brain infusion of S protein in mice induces late cognitive impairment and increases serum levels of neurofilament light chain (NFL), which recapitulates post-COVID features. Neuroinflammation, hippocampal microgliosis and synapse loss are induced by S protein. Increased engulfment of hippocampal presynaptic terminals late after S protein brain infusion were found to temporally correlate with cognitive deficit in mice. Blockage of TLR4 signaling prevented S-associated detrimental effects on synapse and memory loss. In a cohort of 86 patients recovered from mild COVID-19, genotype GG TLR4 -2604G>A (rs10759931) was associated with poor cognitive outcome. Collectively, these findings indicate that S protein directly impacts the brain and suggest that TLR4 is a potential target to prevent post-COVID cognitive dysfunction.

**One Sentence Summary:** TLR4 mediates long-term cognitive impairment in mice and its genetic variant increases the risk of poor cognitive outcome in post-COVID patients.

## INTRODUCTION

Severe acute respiratory syndrome coronavirus 2 (SARS-CoV-2) is considered a respiratory pathogen, but the impact of the infection on extrapulmonary tissues is of high concern. Coronavirus disease 2019 (COVID-19) is associated with unpredictable and variable outcomes, and while most patients show a positive recovery after the acute stages, others experience a myriad of acute and long term neurological dysfunctions*(1)*. Cognitive impairment is a well-characterized feature of the post-COVID syndrome, referred to as “long COVID” or brain fog*(2)*. Mounting evidence suggests that COVID-induced neurological symptoms are mediated by multiple mechanisms, including brain hypoxia and systemic inflammation even in patients with mild symptoms*(3, 4)*. Despite some findings indicating that SARS-CoV-2 can reach and directly impact the brain, others indicate that the virus can rarely cross the blood-brain-barrier (BBB)*(5, 6)*. Nevertheless, whether brain presence of SARS-CoV-2 viral particles and/or its products is a crucial event for the development of cognitive impairment in post-COVID patients remains unknown.

SARS-CoV-2 spike (S) protein plays a pivotal role in COVID-19 pathogenesis and is the main target for vaccine development. This viral surface protein is a homotrimer composed of two functional domains, also known as subunits (S1 and S2), as they are generated by proteolytic cleavage of S protein after virus binding to enzyme 2 angiotensin-converting (ACE2), which mediates cell entry*(7)*. During SARS-CoV-2 infection, cells produce and release variable amounts of viral particles and proteins, including the S protein*(7, 8)*. The S1 was shown to cross the BBB, reaching different memory-related regions of the brain in a mouse model of SARS-CoV-2 infection*(9)*. Inflammation and increased BBB permeability were also shown in *in vitro* models of S1 exposure*(10)*. Likewise, the protein was detected in the central nervous system (CNS) of COVID-19 patients, irrespective of viral RNA detection*(11, 12)*. In addition, increased levels of proinflammatory cytokines and brain gliosis have been reported in severe COVID-19 patients*(13,14)*. Nonetheless, proof concerning the acute and chronic impact of S protein on COVID-19 brain dysfunction and its underlying mechanisms are still lacking.

Most experimental studies investigating the effects of SARS-CoV-2 have focused on acute infection, especially on peripheral tissues. Few studies have used experimental models to evaluate the possible mechanism of post-COVID syndrome. Here, we developed a mouse model of intracerebroventricular (icv) of S exposure to understand the role of this protein in late cognitive impairment after viral infection. Here, we infused S protein in the brains of mice and performed a long-term (45 days) follow-up of the behavioral, neuropathological, and molecular consequences. We report late cognitive impairment, synapse loss, and microglial engulfment of presynaptic terminals after icv infusion of S protein. Early TLR4 blockage prevented S-associated detrimental effects on synapse and memory. We also demonstrated that the single nucleotide polymorphism (SNP) rs10759931, linked with increased TLR4 expression is associated with long-term cognitive impairment in mild COVID-19-recovered patients. Collectively, these findings show that S protein impacts the mouse CNS, independent of virus infection, and identify TLR4 as a key mediator and interesting target to investigate the long-term cognitive dysfunction both in humans and rodents.

## RESULTS

### SARS-CoV-2 spike protein induces long-term cognitive impairment and synapse loss in mice

COVID-19 is associated with long-term cognitive dysfunction*(2)*. To evaluate whether SARS-CoV-2 spike protein induces behavioral changes, we infused the protein into mouse brains through icv route. The experiments were performed in two different timeframes: “early and “late” phases, corresponding to assessments performed within the first 7 days and between 30 and 45 days after S protein infusion, respectively (Fig. 1A). Animals were submitted to behavioral tests or culled to collect brain or blood samples at different time points after infusion (Fig. 1A). To evaluate the effect of S protein infusion on declarative memory, mice were evaluated using the novel object recognition (NOR) test*(15)*. Vehicle-infused mice (Veh) were able to perform the NOR task as demonstrated by a longer exploration of the novel object over the familiar one (Fig. 1B-D, white bars). The S protein had no impact on memory function in the early phase after brain infusion (Fig. 1B, gray bars), while in later time points infused mice failed to recognize the novel object (Fig. 1C-D, black bars). In order to rule out the possibility that changes in motivation, motor function and/or anxiety levels eventually induced by S protein infusion were influencing NOR interpretation, mice were submitted to the rotarod and open field tests. Both S protein-and Veh-infused groups showed similar innate preferences for the objects in the NOR memory test (Supplementary Fig. 1A-C), similar motivation towards object exploration in the NOR sessions (Supplementary Fig. 1D-F) and performed similarly in the rotarod (Fig. 1E) and open field tests (Fig. 1F-G).

**Figure 1.**
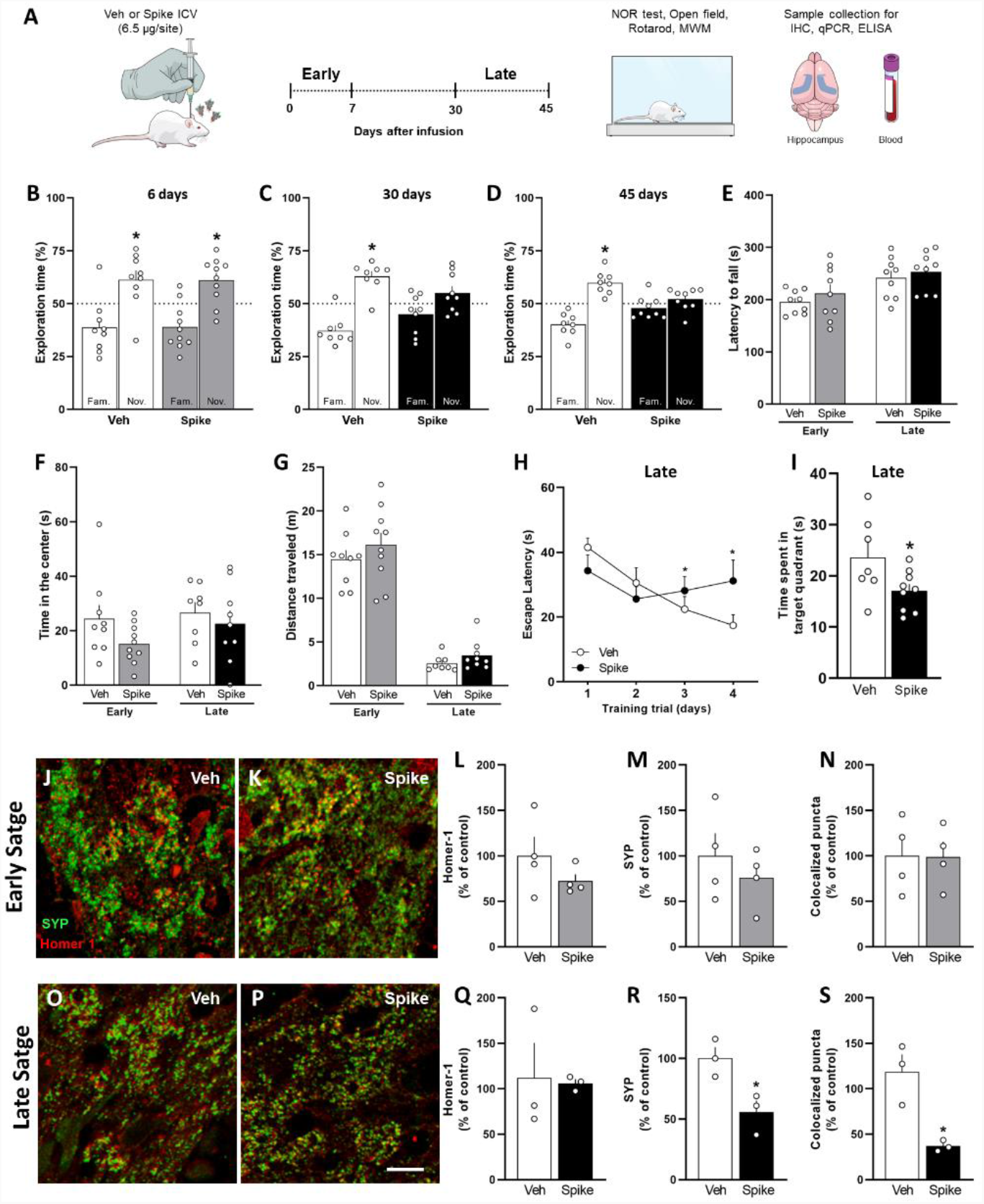
Spike protein causes synapse damage and memory impairment in mice. (**A**) Mice received an i.c.v. infusion of 6,5 μg of SARS-CoV-2 spike protein (Spike), or vehicle (Veh), and were evaluated at early (up to 7 days) or late time points (from 30 to 45 days) after the infusion using behavioral and molecular approaches. (**B to D**) Mice were tested in the novel object recognition (NOR) test at 6 days (**B** t=2.626, \**p* = 0.0304 for Veh; and t=3.218, \**p* = 0.0104, for Spike), 30 days (**C** t=5.099, \**p* =0.0014 for Veh), and 45 days after protein infusion (**D** t=5,122, \**p* = 0.0014, for Veh); one-sample Student’s *t*-test compared to the chance level of 50% (*N* = 8-10 mice per group). (**E**) Mice were tested in the Rotarod task at early and late (*N* = 9 mice per group). (**F**)Time spent at the center of the open field arena at early or late stages of the model (*N* = 8-10 mice per group). (**G**) Total distance traveled of the open field arena at early or late (*N* = 8-10 mice per group). (**H to I**)Escape latencies across 4 consecutive training trials (**H**) and time spent in the target quadrant during the probe trial (I) of the MWM test performed at the late stage (**H** *F*_(3, 45)_ = 2.857, \**p* = 0.0475, repeated measures ANOVA followed by Tukey’s test; **I** t = 2.211, \**p* = 0.0442, Student’s *t*-test; *N* = 7-9 mice per group). Representative images of the DG hippocampal region of Veh-(**J**,**O**) or Spike-infused mice (**K, P**) in the early (**J, K**) and late (**O, P**) stages of the model, immunolabeled for Homer1 (red) and synaptophysin (SYP; green). (**L to N, G to S**) Number of puncta for Homer-1 (**L, Q**), SYP (**M, R**), and colocalized Homer-1/SYP puncta (**N, S**) in the early (**L to N**) and late (**Q to S**) stages of the model. (**R** t = 3.400, \**p* = 0.0273; **s** t = 4.204, \**p* = 0.0137, Student’s *t*-test; *N* = 3-4 mice per group). Scale bar = 20 μm. Symbols represent individual mice. Bars (**B to G**; **I**; **L to N; Q to S**) or points (**H**) represent means ± SEM. OD: optical density; IHC: immunohistochemistry; NOR: Novel object recognition.

Late cognitive dysfunction induced by S protein infusion was confirmed by the Morris Water Maze (MWM) test, a task widely used to assess spatial memory in rodents*(16)*. Mice infused with S protein showed higher latency time to find the submerged platform in sessions 3 and 4 of MWM training, when compared to control mice (Fig. 1H). Also, S protein-infused mice showed reduced memory retention, as indicated by the decreased time spent by these animals in the target quadrant during the probe trial (Fig. 1I). No difference in the swimming speed (Supplementary Fig. 1G) or distance traveled (Supplementary Fig. 1H) were found between groups during the test session.

Synapse loss is strongly correlated to the cognitive decline observed in neurodegenerative diseases*(17)*. Thus, we next investigated whether S protein induces synapse damage in the mouse hippocampus, a brain region critical for memory consolidation. S protein-infused mice did not show changes in synaptic density at the early stages, as demonstrated by the similar immunostaining for synaptophysin (SYP) and Homer-1 (pre-and postsynaptic markers, respectively) compared with the control group (Fig. 1J-M). Equivalent results were also found for the colocalization of these synaptic markers, which indicates no changes in synaptic density (Fig. 1J-K, N). In contrast, decreased SYP immunostaining and synaptic puncta were observed longer periods after S protein infusion (Fig. 1O-S), indicating that spike-induced cognitive dysfunction (shown in Figs. 1D, H, I) displays temporal correlation with hippocampal synapse damage (Fig. 1O-S). Collectively, these data suggest that a single brain infusion of S protein induces late synaptic loss and cognitive dysfunction, mimicking the post-COVID syndrome *(2, 3)*.

### SARS-CoV-2 spike protein triggers late neuroinflammation in mice

Neurodegeneration associated with viral brain infections can be mediated either by direct neuronal injury or by neuroinflammation*(18)*. To advance in the understanding of the genuine impact of SP on neurons, cultured primary cortical neurons were incubated with S protein for 24 h. Neuron exposure to S protein did not affect neuron morphology (Supplementary Fig. 2A-E), once the percent of pyknotic nuclei (Supplementary Fig. 2C), number of primary neurites (Supplementary Fig. 2D) and intensity of β3-tubulin immunostaining (Supplementary Fig. 2E) were similar for vehicle-and S protein-incubated neurons. Also, S protein incubation also had no effect on the neuronal synaptic density and puncta (Supplementary Fig. 2F-J), suggesting that neurons are not directly affected by S protein.

Microglia is the primary innate immune cell of the brain and plays a critical role in neuroinflammation-induced cognitive dysfunction*(19)*. To further understand the impact of spike protein on microglial activation, mouse microglia BV-2 cell lineages were incubated with S protein for 24h. We found that S protein stimulation increased Iba-1 immunoreactivity (Supplementary Fig. 2K-M) and upregulated TNF, INF-β and IL-6 expression (Supplementary Fig. 2N-P), without affecting IL-1β and IFNAR2 (Supplementary Fig. 2Q-R). To evaluate the time course of the in vivo activation of microglia, we analyzed cellular features and cytokine production in our mouse model. We found that at the early stage after icv. injection of S protein neither changed the number and morphology of microglia (Fig. 2A-D) nor increased the levels of TNF-α, IL-1β, IL-6, INF-β and IFNAR1 (Fig. 2E-I). In contrast, the levels of IFNAR2 mRNA decreased significantly at the same time point after S protein infusion (Fig. 2J). Notably, assessments performed late after revealed an increased number of Iba-1-positive cells (Fig. 2K-M) and a predominance of cells with ameboid morphology in the hippocampus (Fig. 2K, L, N), suggestive of microglial cells in a reactive state. However, no differences in GFAP immunoreactivity were detected in S protein-infused mice when compared to the control group (Supplementary Fig. 3A-C). The level and/or expression of the proinflammatory cytokines TNF, IL-1β, IFNα and IFNβ (Fig. 2O-S) and the receptor IFNAR2 (Fig. 2T) were higher in the hippocampus of S protein brain infused mice at this time point. Hippocampal expression of IL-6 and IFN-γ cytokines and the receptor IFNAR1 were unaffected by S protein infusion (Supplementary Fig. 3D-F). Altogether, our results indicate that the cognitive impairment induced by S protein is accompanied by microglial activation and neuroinflammation.

**Figure 2.**
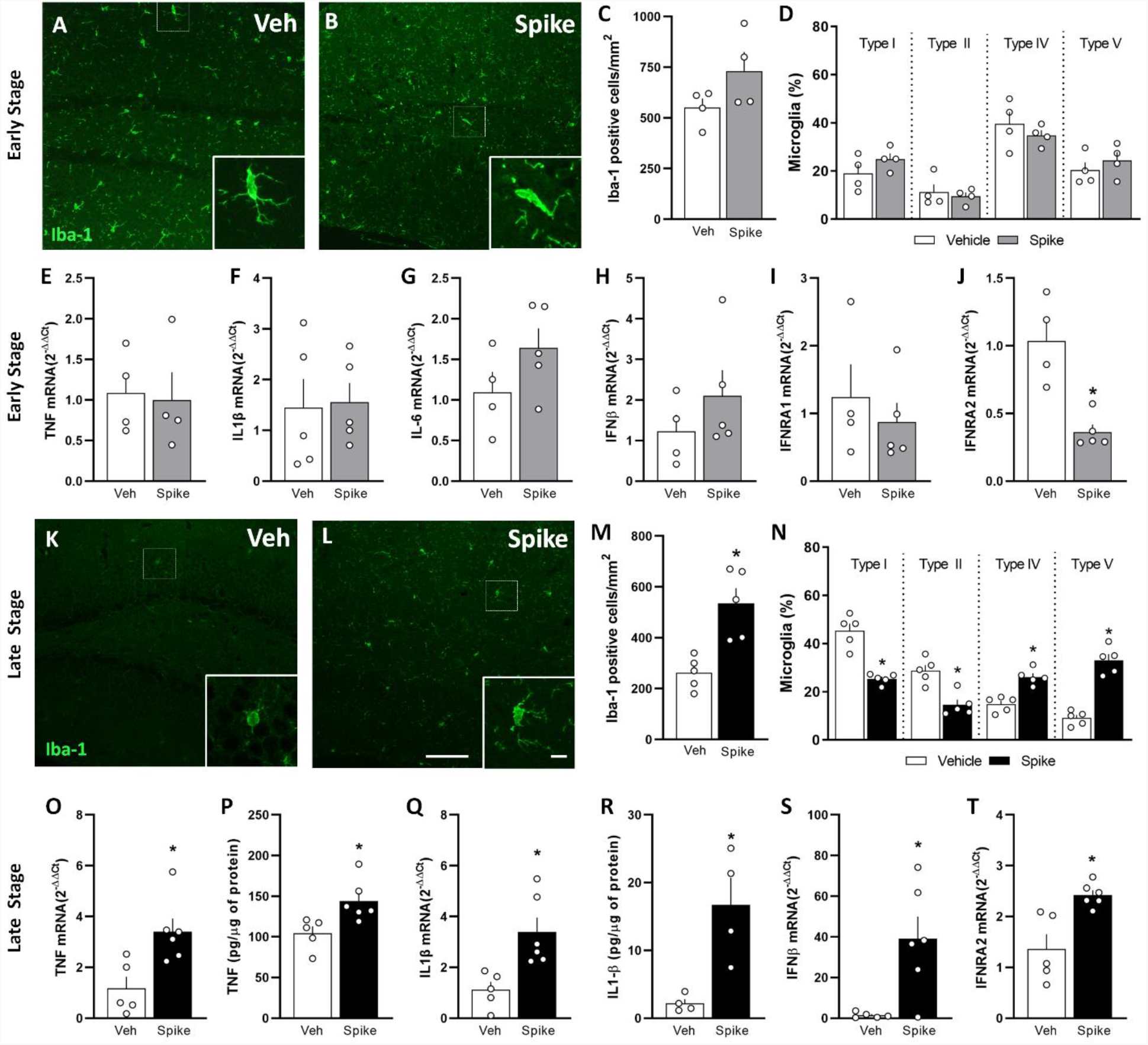
Spike protein induces cytokine upregulation in cultured microglia and triggers delayed brain inflammation and microgliosis in mice. **a-t**, Mice received an i.c.v. infusion of 6,5 μg of Spike or vehicle (Veh), and were evaluated at early (**a-j**, 3 days) or late time points (**k-t**, 45 days). Representative images of Iba-1 immunostaining in the DG hippocampal region of Veh-(**a, k**) or Spike-infused mice (**b, l**) in the early (**a, b**) and late (**k, l**) stages of the model. Scale bar = 25 μm, inset scale bar = 10 μm. **c, m**, Iba-1 positive cells in the hippocampi of Veh-or Spike-infused mice in the early (**c**) and late (**m** t = 4.086; \**p* = 0.0035, Student’s *t*-test) stages of the model (*N* = 4-5 mice per group). **d, n**, Quantifications of the proportion of each Iba-1-positive cells morphological type in Veh-or Spike-infused mice evaluated in the in the early and late (**n** t = 6.388; \**p*= 0.0002 for Type I; t = 4.458; \**p* = 0.0021 for Type II; t =5.513; \**p* =0.0006 for Type IV; t = 8.384; \**p* < 0.0001 for Type V, Student’s *t*-test) stages of the model (*N* = 4-5 mice per group). **e-j**, qPCR analysis of indicated mRNA isolated from the hippocampus in the early stage of the model. IFNRA2 (**j** t=4.413, \**p* = 0.0031) (*N* = 4-5 mice per group). **o-t**, Hippocampal proinflammatory mediators in Veh-or Spike-infused mice in the late stage of the model. TNF mRNA (**o** t=3.189; \**p* = 0.0110) and protein (**p** t=2.885; \**p* =0.0180) levels. IL-1β mRNA (**q** t=3.322; \**p* = 0.0089) and protein (**r** t=3.583; \**p* =0.0116) levels. **s-t** mRNA levels of IFN-β (**s** t=3.713, \**p* =0.013) and IFNAR2 (**t** t=3.743; \**p* = 0.0046). Student’s *t*-test (*N* = 4-6 mice per group). Symbols represent individual mice, and bars represent means ± SEM.

**Figure 3:**
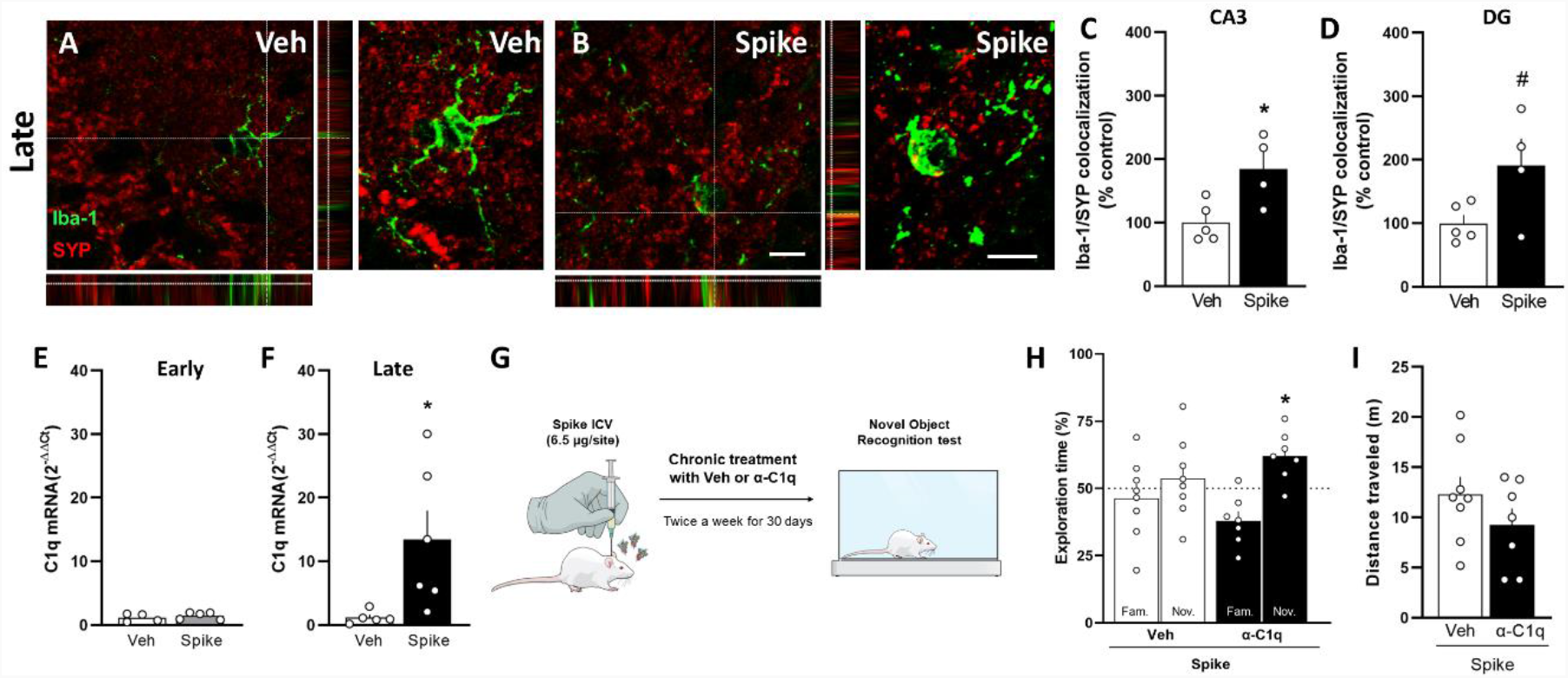
C1q neutralization prevents spike-induced memory impairment in mice. Mice received an i.c.v. infusion of 6,5 μg of SARS-CoV-2 spike protein (Spike), or vehicle (Veh), and were evaluated at early (3 days) or late time points (45 days). (**A to B**) Representative images of microglia (Iba-1^+^, green) engulfing pre-synaptic terminals immunolabeled for synaptophysin (SYP, red) in the DG hippocampal region of Veh-(**A**) or Spike-infused mice (**B**) in the late stage of the model. Scale bar = 25 μm, inset scale bar = 10 μm. (**C to D**) Quantification of microglia-SYP colocalization in CA3 (**C** t = 2.949, \**p* = 0.0214), and DG (**D** t = 2.271, ^#^*p* = 0.0574) hippocampal regions. Student’s *t*-test; (*N* = 4-5 mice per group). (**E to F**) C1q mRNA expression in hippocampi of Veh-or Spike-infused mice at early (**E**) or late time points (**F** t = 2.425, \**p* = 0.0383, Student’s *t*-test; (*N* = 4-6 mice per group). (**G**) Mice received an i.c.v. infusion of 6,5 μg of Spike, were treated with Veh or 0.3 μg anti-C1q antibody (α-C1q; i.c.v., twice a week, for 30 days), followed by novel object recognition (NOR) testing (**H** t=3.438, \**p* = 0.0138 for Spike/α-C1q; one-sample Student’s *t*-test compared to the chance level of 50%). (**I**) Total distance traveled of the open field arena at late (*N* = 7-8 mice per group).

### SARS-CoV-2 spike protein induces C1q-mediated synaptic phagocytosis by microglia in mice

Synaptic phagocytosis (or synaptic pruning) by microglia was shown to underlie cognitive dysfunction in dementia and in viral encephalitis*(17, 20)*. We therefore evaluated whether synaptic phagocytosis by microglia mediates S protein-induced synapse damage. Hippocampal three-dimensional image reconstructions of Iba-1-positive cells from S protein-infused mice showed increased SYP-positive terminals inside phagocytic cells (Fig. 3A-D). The complement component 1q (C1q) protein is known to be involved in the initial tagging of synapses, preceding synaptic engulfment by microglial cells*(21)*. Accordingly, we found that C1q was significantly upregulated in the brains of mice late (but not early) after S protein infusion (Fig. 3E, F). This finding led us to investigate whether or not the blockage of soluble C1q, using a neutralizing antibody, could restore cognitive function in S protein-infused mice. For this, the animals were treated by icv. route with a neutralizing C1q antibody immediately after S protein infusion and twice a week for 30 days (Fig. 3G). Remarkably, C1q blockage rescued memory impairment in S protein-infused mice (Fig. 3H), without any effect on locomotion (Fig. 3I) or exploration (Supplementary Fig. 4A, B). As seen for many viral encephalitis, these data suggest that C1q-mediated microglial phagocytosis underlie long-term cognitive dysfunction induced by S protein.

### TLR-4 mediates cognitive dysfunction induced by SARS-CoV-2 spike protein

Recent findings have described that S protein induces toll-like receptor 4 (TLR4) activation in cultured immune cells*(22)*. Additionally, TLR4 has been implicated in microglial activation and cognitive dysfunction in degenerative chronic disease of CNS like Alzheimer’s disease*(23)*. In agreement with these observations, despite no changes found in TLR4 expression levels at the early time point after S protein infusion (Fig. 4A), we found a late upregulation of TLR4 gene (Fig. 4B) in the hippocampus of infused mice that matches the late cognitive dysfunction (shown in Figs. 1C-D, H-I). To evaluate the role of TLR4 in spike-induced cognitive impairment we used either a pharmacological approach or a TLR4 knockout mouse model (TLR4^-/-^). First, to investigate whether activation of TLR4 is an early event that could impact cognition later on, mice were treated with the TLR4 inhibitor TAK242 1h before S protein brain infusion and once a day for 7 days (Fig. 4C). Remarkably, early inhibition of TLR4 greatly prevented late memory dysfunction induced by S protein infusion in mice (Fig. 4D). Recent evidence has shown that high plasmatic levels of Neurofilament-light chain (NFL) are correlated with poor outcome in and COVID-19 patients*(6, 14, 24)*. Then, we evaluated the NFL levels in plasma samples of control and S protein-infused mice, treated or not with TAK242 (Fig. 4E). Using transgenic mice, in the early phase after infusion, both WT and TLR4^-/-^ mice learned the NOR task (Supplementary Fig. 4C). On the other hand, when evaluated at a late time point after protein infusion, WT mice had a poor performance in NOR test, while TLR4^-/-^ animals were able to execute the task (Fig. 4F). Also, the absence of TLR4-mediated response in the TLR4^-/-^ mice prevented the reduction of SYP-positive terminals inside phagocytic cells later after S protein infusion in comparison to WT mice (Fig. 4G-K). Consistent with the previous results, control experiments showed that genetic (Supplementary Fig. 4D-I) or pharmacological (Supplementary Fig. 4J-L) inhibition of TLR4 had no effect on locomotion or exploratory behavior. Finally, we also found reduced microgliosis (Fig. 4L-O) and less microglia-engulfed synapses in the hippocampus of TLR4-/-mice later after S protein brain infusion (Fig. 4P-S). Together, these data suggest that TLR4 activation mediates cognitive deficit and synaptic pruning induced by S protein in mice.

**Figure 4:**
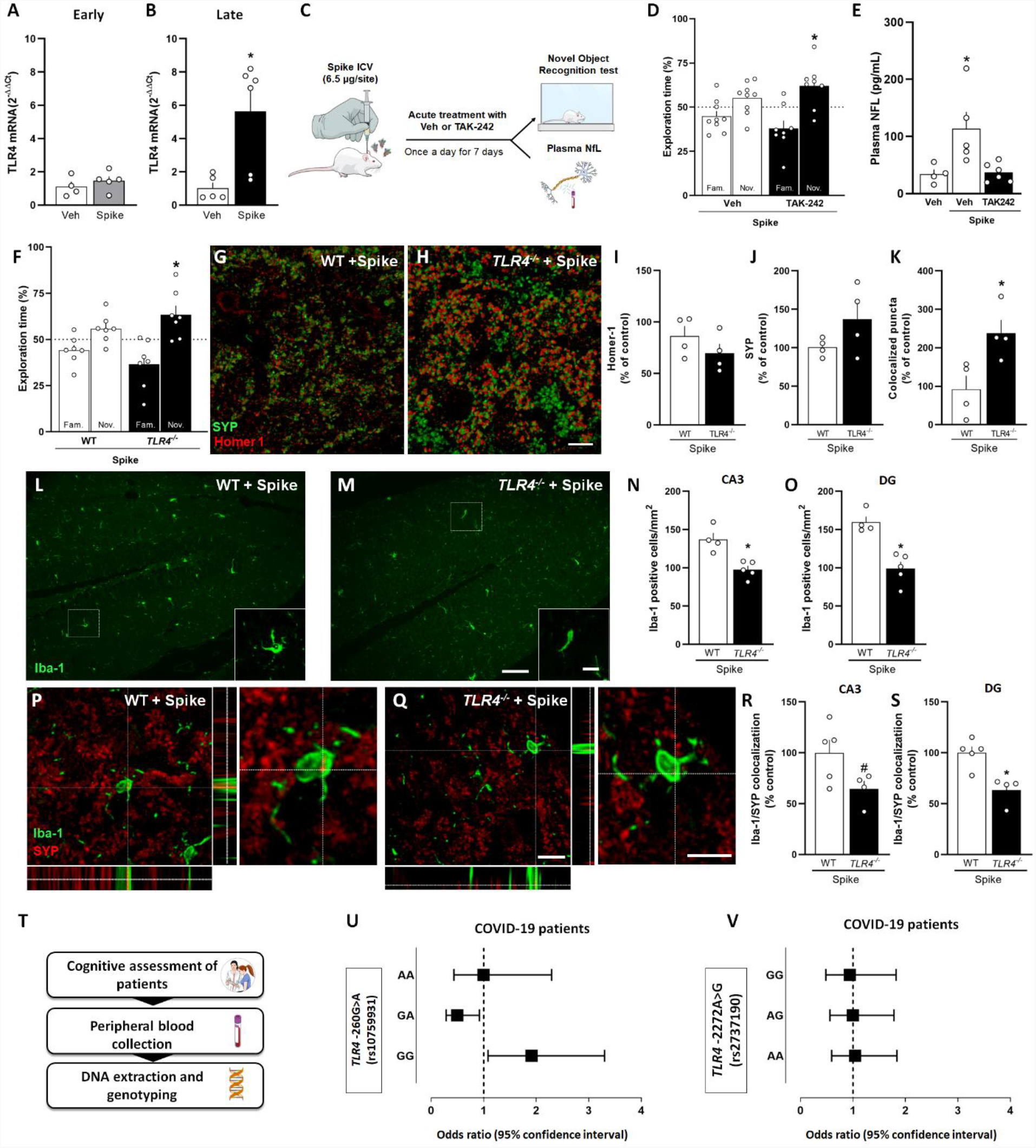
TLR4 mediates spike-induced memory impairment in mice and is associated with post-COVID cognitive impairment in a human cohort. **a, b**, Mice received an i.c.v. infusion of 6,5 μg of SARS-CoV-2 spike protein (Spike), or vehicle (Veh), and TLR4 mRNA levels in the hippocampi of Veh-or Spike-infused mice were evaluated at early (**a**; 3 days) or late (**b**, 45 days, t = 3.229, \**p* = 0.0103, Student’s *t*-test) time points (*N* = 4-6 mice per group). **c**, Swiss mice received an i.c.v. infusion of 6,5 μg of Spike and were treated with Veh or the TLR4 antagonist TAK-242 (2 mg/kg, once daily for 7 days, i.p.), and were tested in the late stage of the model in the novel object recognition (NOR) test (**d** t = 2.713, \**p* = 0.0301 for Spike/TAK-242; one-sample Student’s *t*-test compared to the chance level of 50%, *N* = 8-9 mice per group). **e** Plasma NfL levels evaluated in the late stage of the Spike infusion model (F = 6.329, \**p* = 0.0133, one-way ANOVA test, followed by Tukey’s test (*N* = 4-6 mice per group). **f**, Wild-type (WT) and TLR4 knockout (*TLR4*^*-/-*^) mice received an icv infusion of 6,5 μg of SARS-CoV-2 spike protein (Spike) and were tested in the novel object recognition (NOR) test in the late stage of the model (**f** t = 2.744, \**p* = 0.0336). One-sample Student’s *t*-test compared to the chance level of 50%, *N* = 7 mice per group). **g, h**, Representative images of the DG hippocampal region of WT/Spike (**g**) or *TLR4*^*-/-*^*/*Spike (**h**) mice immunolabeled for Homer1 (red) and synaptophysin (SYP; green). Scale bar = 20 μm. Number of puncta for Homer-1 (**i**), SYP (**j**), and colocalized Homer-1/SYP puncta (**k** t = 2.945, \**p* = 0.0258; Student’s *t*-test; *N* = 4 mice per group). **l, m**, Representative images of Iba-1 immunolabeling in the DG hippocampal region of WT (**l**) or *TLR4*^*-/-*^ (**m**) mice infused with Spike. Scale bar = 25 μm, inset scale bar = 10 μm. Iba-1 positive cells in the CA3 (**n** t = 4.242; \**p* = 0.0038) and DG (**o** t = 5.088; \**p* = 0.0014) hippocampal regions of WT or *TLR4*^*-/-*^ mice infused with Spike. **p, q**, Representative images of microglia (Iba-1^+^, green) engulfing pre-synaptic terminals immunolabeled for synaptophysin (SYP, red) in the DG hippocampal region of WT (**p**) or *TLR4*^*-/-*^ (**q**) mice infused with Spike. Scale bar = 50 μm, inset scale bar = 10 μm. **r, s**, Quantification of microglia-SYP colocalization in CA3 (**r** t = 2.200, ^#^*p* = 0.0637), and DG (**s** t = 4.012, \**p* = 0.0051) hippocampal regions. Student’s *t*-test; (*N* = 4-5 mice per group). Symbols represent individual mice, and bars represent means ± SEM. **t**, Pipeline to analyze the impact of *TLR4* variants in cognitive status of Post-COVID patients. **u, v**, Forest plots showing odds ratio and 95% confidence interval for risk of cognitive impairment post-COVID-19 by genotype for SNPs *TLR4* - 2604G>A (rs10759931; **u**) and *TLR4* − 2272A>G (rs2737190; **v**). Each square represents the odds ratio for each genotype, and each horizontal line shows the 95% confidence interval.

Importantly, the early treatment with TLR4 inhibitor prevented the late neuronal damage, indicating that the TLR4 pathway is central to induce neurodegeneration and long-term cognitive impairment in the present model.

### Single nucleotide polymorphism within TLR4 gene is associated with increased risk of cognitive dysfunction after COVID-19

Several lines of evidence have suggested that polymorphisms in TLR4 is a risk factor for developing inflammatory diseases, including sporadic Alzheimer’s disease*(23, 25)*. Thus, we sought to extend our findings by investigating whether there is an association between TLR4 gene variants and cognitive outcomes in COVID-19 patients. For this, 86 individuals with confirmed COVID-19 diagnosis, mostly with mild disease, were included in the study sample (Fig. 4L). Characteristics of the sample are displayed in Supplementary Table 1. Cognition was assessed using the Symbol Digit Modalities Test (SDMT) from 1 to 16 months after the onset of COVID-19 acute symptoms (Mean: 6.85 months). Of interest, nearly half of the patients evaluated (40; 46.51%) presented an important degree of post-COVID-19 cognitive impairment (Table 1). Genotyping analysis for two different SNPs (rs10759931 and rs2737190) was performed in all studied subjects. Individuals carrying the *TLR4*-2604G>A (rs10759931) GG homozygous genotype demonstrated a significantly higher risk for developing cognitive impairment following SARS-CoV-2 infection (*p*-value = 0.0234; OR= 1.91), while the GA genotype was associated with a decreased risk (*p*-value = 0.0209; OR= 0.50) (Fig. 4M and Table 1). Conversely, none of the *TLR4-2272A>G* (rs2737190) genotype variations were associated with increased susceptibility to post-COVID-19 cognitive impairments (Fig.4N and Table 1). We then hypothesized that polymorphisms in TLR4 gene are probably associated with altered spike-induced host immune responses, increasing the risk to develop long-term cognitive deficit in genetically susceptible individuals.

**Table 1.** *TLR4* rs10759931 and rs2737190 genotype distribution in patients with or without cognitive deficit following COVID-19.

| **$TLR4 - 2604\text{G}>\text{A}$**<br>**(rs10759931)** | **N**<br>**(86)** | **Cognitive**<br>**Deficit**<br>**(%)** | **No**<br>**Cognitive**<br>**Deficit (%)** | **P-**<br>**value** | **OR (95% CI)** |
| --- | --- | --- | --- | --- | --- |
| GG | 40 | 22(55) | 18(39) | 0.0234<br>* | 1.91 (1.083 to<br>3.301) |
| GA | 35 | 13(32) | 22(48) | 0.0209<br>* | 0.50 (0.287 to<br>0.920) |
| AA | 11 | 5(13) | 6(13) | >0,999<br>9 | 1.00 (0.435 to<br>2.294) |
| MAF (A) | 0.33 |  |  |  |  |
| **$TLR4 - 2272\text{A}>\text{G}$**<br>**(rs2737190)** | **N**<br>**(83)** | **Cognitive**<br>**Deficit**<br>**(%)** | **No**<br>**Cognitive**<br>**Deficit (%)** | **P-**<br>**value** | **OR (95% CI)** |
| AA | 30 | 14(37) | 16(36) | 0.8832 | 1.04 (0.594 to 1.836) |
| AG | 35 | 16(42) | 19(42) | >0,999<br>9 | 1.0 (0.561 to 1.781) |
| GG | 18 | 8(21) | 10(22) | 0.8633 | 0.94 (0.483 to 1.823) |
| MAF (G) | 0.43 |  |  |  |  |

MAF= minor allele frequency; OR = odds ratio; CI = confidence interval. Data analyzed by $\chi^2$ -test (two-tailed). \*Statistical significance ( $P<0.05$ ). The reference group in each of the analyses was the most prevalent genotype.

## DISCUSSION

Long COVID comprises a myriad of symptoms that emerge after the acute phase of infection, including psychiatric symptoms, and dementia-like cognitive dysfunction *(1, 3, 4, 26– 28)*. Clinical studies have largely mapped the spectrum of neurological symptoms in post-COVID patients, but do not provide significant advance in describing the molecular mechanisms that trigger this condition or targets for preventive/therapeutic interventions. In contrast, studies involving COVID-19 preclinical models have entirely focused on the acute impacts of viral infection. Therefore, it is mandatory to develop novel tools to dissect the mechanisms underlying the neurological deficits in long COVID, especially the direct effect of the virus and/or viral products on the brain.

It has been suggested that the S protein can be released from virions*(29, 30)*, suggesting that it could directly trigger brain damage. Thus, we speculated that S plays a central role in neurological dysfunctions associated with COVID-19, independently of SARS-CoV-2 replication in the brain. Previous studies demonstrated that the hippocampus is particularly vulnerable to viral infections*(15, 20, 31)*. Accordingly, brain scans of COVID-19-recovered patients showed significant changes in hippocampal volume*(32)*, an important predictor of cognitive dysfunction in both normal aging and Alzheimer’s disease *(33, 34)*. Here, we developed a rodent model that mimics key neurological features of long COVID through brain icv infusion of S. Using two hippocampal-dependent behavioral paradigms, we found that brain exposure to S disrupts long-term mouse memory, with no early behavioral impact. To our knowledge, this animal model is the first to recapitulate the late cognitive impact of COVID-19.

Synapse damage is a common denominator in a number of memory-related diseases*(35)*, often preceding neurodegeneration. It has been shown that neuroinvasive viruses, such as West Nile virus (WNV), Borna disease virus (BDV) and Zika virus (ZIKV), are also associated with synapse impairment*(15, 20, 36)*. Likewise, we found that the late cognitive dysfunction induced by S was accompanied by prominent synapse loss in mice hippocampus. Recent data have revealed the upregulation of genes linked to synapse elimination in SARS-CoV-2-infected human brain organoids and in post-mortem samples from COVID-19 patients*(11, 37)*. In line, we found that infusion of S into mouse brain induces a late elevation in plasma levels of NFL, an axonal cytoskeleton protein recently identified as a component of pre-and postsynaptic terminals*(38)*. Plasma NFL increase can be employed as a marker of synapse loss and disease progression in neurodegenerative diseases, including Alzheimer’s disease*(39)*. Remarkably, recent data showed that plasma NFL levels are higher in patients with severe COVID-19 compared to healthy age-matched individuals, as well as inversely correlated to the cognitive performance in COVID-19 patients*(40, 41)*, reinforcing the translational potential of our model. Collectively, these findings suggest that brain exposure to S induces the synapse loss and behavioral alterations typical of viral encephalitis, leading to a prolonged neurological dysfunction that can persist long after recovery from the infectious event.

Microglia are the most abundant immune cell type within the CNS and play a critical role in most of the neuroinflammatory diseases *(42)*. In viral encephalitis, microglial cells have both protective and detrimental activities depending on the phase of infection*(19)*. Previous studies showed that human coronaviruses can reach the CNS and induce gliosis both in mature and immature brain tissues *(12, 31, 43)*. Here we found that microglial cell lineage BV-2 was impacted by S protein, corroborating recent data showing an increase in proinflammatory mediators in S1-stimulated microglia*(44)*. Since cultured primary cortical neurons were not directly affected by S stimulation, our *in vitro* results indicate that microglia could be seen as the main cell type affected by exposure to SARS-CoV-2 S protein.

It is well known that viral infections are often associated with excessive activation of inflammatory and immune responses, which may in turn elicit and/or accelerate brain neurodegeneration*(45)*. Here, we found that S-infused mice presented late microglial activation, but not astrocyte reactivity, similar to observed in other animal models of viral encephalitis*(15, 20)*. Hippocampal increased levels of proinflammatory mediators were found only at late time points after S infusion, showing a temporal correlation with synaptic loss and cognitive dysfunctions. Conversely, we found that the downregulation of *IFNAR2* gene occurred shortly after S injection, similar to what is observed in neuronal cells of post-mortem COVID-19 patients*(11)*. This finding corroborates recent evidence demonstrating that SARS-CoV-2 may evade innate immune through modulation of type-I IFN responses *(46)*. Altogether, our results show that brain exposure to S induces an early negative modulation of the main receptor involved in type-I IFN response followed by a late proinflammatory process in the hippocampus.

A complement-microglial axis has emerged as one of the key triggers of synapse loss in memory-related diseases*(17, 21)*. The classical complement cascade, a central player of innate immune pathogen defense, orchestrates synaptic pruning by microglia during physiological and pathological conditions*(47, 48)*. We have previously reported that hippocampal synapses are phagocytosed by microglia during ZIKV brain infection, in a process dependent on C1q and C3*(15)*. Moreover, Vasek and colleagues (2016) showed hippocampal synapse loss in post-mortem samples of patients with WNV neuroinvasive disease, as well as complement-dependent microglial synapse engulfment in both WNV-infected and -recovered mice*(20)*. Accordingly, we demonstrated that cognitive impairment induced by S protein is associated with hippocampal C1q upregulation and microglial engulfment of presynaptic terminals. Additionally, chronic C1q neutralization preserved memory function in S-infused mice, supporting the role of C1q-mediated synaptic pruning as an important mediator of long COVID cognitive impairment.

The pattern recognition receptor TLR4 has been implicated in the neuropathology of viral encephalitis classically associated with memory impairment, including those caused by WNV, Japanese encephalitis virus (JEV) and BDV, as well as age-related neurodegenerative diseases*(49– 52)*. Notably, *in silico* simulations predicted that the S protein could be recognized by the TLR4*(53, 54)*, with this interaction activating the inflammatory signaling, independently of ACE2*(22, 44, 55, 56)*. Accordingly, here we found that a single brain infusion of S protein induced hippocampal TLR4 upregulation. To gain further insight into the role played by TLR4 in COVID-19-induced brain dysfunction, we first performed the pharmacological blockage of TLR4 signal transduction early after S protein brain infusion. This strategy significantly prevented the long-term cognitive impairment observed in our model. Likewise, late cognitive impairment induced by S protein was absent in TLR4-deficient mice, in accordance with previous findings in animal models of dementia*(57, 58)*. Remarkably, we also found that S-induced plasma NFL increase was dependent on TLR4 activation, as early TLR4 inhibition mitigated changes in NFL levels. Together, our findings strongly suggest that brain dysfunction in post-COVID is associated to S-induced TLR4 signaling in microglial cells*(59, 60)*.

The engagement of complement and TLRs in signaling crosstalk has been proposed to regulate immune and inflammatory responses in neurodegenerative diseases*(61)*. Indeed, it was shown that TLR4 activation induces the upregulation of complement components in the mouse hippocampus *(62, 63)*. Given the role of complement activation in synaptic pruning, we hypothesized that TLR4 is the molecular switch that regulates microglial synaptic engulfment. Our data showed that absence of TLR4 confers protection against S-induced microglial mediated synaptic pruning, reinforcing the notion that aberrant immunity activation disrupts synaptic integrity and leads to cognitive dysfunction following pathogenic insult.

Finally, and relevantly, we validated our preclinical findings by examining whether TLR4 genetic variants could be associated with poor cognitive outcome in COVID-19 patients with mild disease. In a cohort of mild COVID-19 patients carrying the GG genotype of *TLR4* -2604G>A (rs10759931) variant, we identified a significant association between this genotype and the risk for cognitive impairment after SARS-CoV-2 infection. The G allele has already been associated with increased risk for different disorders with immunological basis, including cardiovascular diseases*(64)*, diabetes-associated retinopathy*(65)*, cancer*(66)*, and asthma*(67)*. On the other hand, the A allele can affect the binding affinity of the TLR4 promoter to transcription factors, culminating in lower expression of this gene in the allele carriers*(68)*. Taken together, our findings suggest that the complex crosstalk between TLR4, complement system and neuroinflammation are important events that determines the development of neurological symptoms in long COVID patients.

There are some limitations to this study. First, despite experimental evidence demonstrating that spike crosses the BBB, there is no available data on the amount of spike that reaches the CNS in the course of SARS-CoV-2 infection, and additional studies are needed to establish the dose-dependent effects of spike administration. This study would benefit from additional approaches to set up new animal models of long COVID. Second, although our study holds translational potential, our findings are limited by the number of patients and SNPs evaluated, and the absence of longitudinal assessments. Extending these investigations to a larger group of patients, with varying degree of cognitive impairment, would allow to precise these first findings.

The impact of long COVID emerges as a major public health concern, due to the high prevalence of prolonged neurological symptoms among survivors. Therefore, strategies designed to prevent or treat neurological long COVID symptoms constitute an unmet clinical need. Our study described a new animal model that recapitulates the long-term impact of the exposure to SARS-CoV-2 S protein on cognitive function. We found that S-induced cognitive impairment triggers innate immunity activation through TLR4, culminating with microgliosis, neuroinflammation and synaptic pruning. The translational value of our model is supported by the correlation between increased plasma NFL and behavioral deficits, as well as by the association between TLR4 genetic status and SARS-CoV-2 cognitive outcomes of recovered COVID-19 patients. Altogether, our findings open new avenues for the establishment of interventional strategies towards prevention and/or treatment of the long-term brain outcomes of COVID-19.

## MATERIALS AND METHODS

### Study Design

The objective of this study was to assess the direct impact of SARS-CoV-2 S protein on cognitive function, and to gain insight into the mechanisms underlying COVID-19 long-term brain effects, thereby identifying new avenues for therapeutic intervention. Dissection of the underlying mechanisms was performed using qPCR, ELISA, SIMOA, behavioral, and immunohistochemical approaches in mice subjected to pharmacological inhibition of C1q or TLR4, with the role of TLR4 being confirmed using a knockout mouse line. Additionally, the direct impact of spike in different brain cell types was assessed using murine BV-2, and primary neuronal cortical cell cultures. Finally, to support the translational relevance of our results, we investigated two *TLR4* SNPs in a cohort of recovered COVID-19 patients with cognitive impairment*(15, 69–71)*. Further details of the study are provided in the corresponding sections of the Supplementary Materials.

### Data analysis

The software Prism v8 (GraphPad) was used for all statistical tests, and values of *p* ≤ 0.05 were considered statistically significant. Student’s *t*-test was applied to analyze qPCR, ELISA, and immunohistochemical data. For NOR experiments, data were analyzed using a one-sample Student’s *t*-test compared to a fixed value of 50%. MWM and NFL measurements were analyzed using repeated measures or one-way ANOVA followed by Tukey’s test, respectively. Allelic frequencies were determined by direct count of the alleles. Genotypic distributions in Hardy– Weinberg equilibrium were evaluated by two-tailed χ^2^-test. The significant differences in allelic and genotypic frequencies were evaluated by Fisher’s exact test and two-tailed χ^2^-test.

## List of Supplementary Materials

Materials and Methods

Fig. S1 Controls of behavioral tests and serological analyses. Fig. S2 Microglial and neuronal culture.

Fig. S3 GFAP immunohistochemistry and qPCR analysis. Fig. S4 Controls of behavioral tests.

Table S1. Participant demographics of the study sample. Table S2: List of primers used in qPCR analyses.

## Acknowledgments

The authors thank Luis F. Fragoso and Ana Claudia Rangel for their technical support; Prof. Leda R. Castilho and her team at Cell Culture Engineering Laboratory (LECC) of COPPE/UFRJ for providing the trimeric spike protein in the prefusion conformation; and Prof. João B. Calixto (Centro de Inovação e Ensaios Pré-clínicos-CIENP), Prof. João B. Teixeira da Rocha (Federal University of Santa Maria) and Prof. Andreza F. De Bem (Brazilian Federal University) for thoughtful discussion and manuscript critical reading.

## Author contributions

C.P.F., G.F.P., S.V.A.L., A.T.P., J.R.C., R.C., and F.L.F.D. conceived the study. C.P.F., G.F.P., S.V.A.L., A.T.P., F.L.F.D., J.R.C., R.C., J.L.S., and L.E.B.S., contributed to experimental design. F.L.F.D., G.G.F., E.V.L., L.S.A., H.P.M.A., L.C.C., S.M.B.A., and T.N.S. performed experiments in mice and analyzed the data. Molecular experiments and ELISA were performed and analyzed by F.L.F.D., L.C.C., S.M.B.A., and A.T.P. Histological and immunohisto/cytochemistry analyses were performed by G.G.F., C.P.F., and E.V.L. L.E.B.S, L.R., and G.G.F. performed experiments in cell culture. L.A.D. and A.L.S. performed Simoa experiments. E.G.G., M.B.H., K.L.P., C.C.F.V., and S.V.A.L. recruited patients, collected clinical information and performed neuropsychological evaluations. L.A.A.L. performed molecular and serological diagnosis of COVID-19. F.L.F.D. and E.G.G. carried out genotype analyses. F.L.F.D., G.G.F., E.G.G., C.P.F., G.F.P., S.V.A.L., A.T.P., J.R.C., R.C., and L.E.B.S. contributed to critical analysis of the data. F.L.F.D., C.P.F, and G.F.P. wrote the manuscript. All authors read and approved the final version.

## Data and materials availability

All data are available in the main text or the supplementary materials.

