## Supplementary materials for "SARS-CoV-2 spike protein induces TLR-4-mediated long-term cognitive dysfunction recapitulating post-COVID syndrome"

**SARS-CoV-2 spike protein induces synaptic loss and cognitive impairment mediated by TLR4**

Fontes-Dantas F.L. et al.

* Corresponding author: A.T.D.P., S.V.A.L, G.P.F. or to C.P.F..

**The PDF file includes:**

Materials and Methods

Fig. S1, S2, S3 and S4

Table S1 and S2

References for Materials and Methods

**Materials and Methods**

**Animals**

Eight to twelve-week-old male Swiss were used in this study. In some experiments, TLR4^-/-^ mice on the C57BL/6 background were used. Animals were housed in groups of five per cage with free access to food and water, under a 12 h light/dark cycle, with controlled temperature and humidity. All procedures followed the “Principles of Laboratory Animal Care” (US National Institutes of Health) and were approved by the Institutional Animal Care and Use Committee of the Federal University of Rio de Janeiro (protocol number 068/2).

**Spike intracerebroventricular infusion**

The recombinant trimeric SARS-CoV-2 spike protein (S; 1-1208aa) in the prefusion conformation produced in HEK293 cells was obtained from Cell Culture Engineering Laboratory (LECC) of COPPE/UFRJ and was produced as described elsewhere. For intracerebroventricular (icv.) infusion of the S protein, mice were anesthetized with 2.5% isoflurane (Cristália; São Paulo, Brazil) using a vaporizer system (Norwell, MA), and a 2.5 mm-long needle was unilaterally inserted 1 mm to the right of the midline point equidistant from each eye and parallel to a line drawn through the anterior base of the eye (REF). Using a Hamilton syringe, 6.5 µg S protein or vehicle (PBS; 137 mM NaCl, 10 mM sodium phosphate, 2.7 mM KCl, pH 7.4) were slowly infused. In order to assess antibody production following S administration, mice received 10 µg S protein or vehicle (PBS) subcutaneously (sc), and boosted after 15 days. The trials were divided into two distinct stages: early phase (assessments performed up to one week after administration) and late phase (between 30 and 45 days after administration).

**Pharmacological treatments**

For TLR4 blockade, TAK-242 (Millipore) was diluted in sterile saline (vehicle) and injected intraperitoneally (ip; 2mg/kg). Mice received either vehicle or TAK for 7 days beginning immediately after S protein icv. administration. For brain C1q blockade, mice received icv. injections of vehicle (PBS) or an antibody against C1q (0.3 μg; Abcam #11861) twice a week for 30 days after S brain infusion.

**Behavioral tests**

*Open field test:* Animals were placed in the center of an arena (30 × 30 × 45 cm) divided in nine imaginary quadrants, and exploration was assessed for 5 min. The arena was thoroughly cleaned with 70% ethanol in between trials to eliminate olfactory cues. Total locomotor activity and time spent at central or peripheral quadrants were analyzed using ANY-maze software (Stoelting Company).

*Novel object recognition (NOR) test:* The test was carried out in an arena measuring 30 × 30 × 45 cm. Before training, each animal was submitted to a 5-min habituation session in the empty arena. Test objects were made of plastic and had different shapes, colors, sizes, and textures. Innate object preferences or neophobia were excluded in preliminary tests. Mice explored the configuration of two identical objects during a 5-min acquisition trial. After 90 min, mice were submitted to a 5-min retention trial, during which one of the familiar objects was replaced by an unfamiliar new one. Sniffing and touching the object were considered exploratory behavior. Results were expressed as a percentage of time exploring each object during the training or test sessions, or as total exploration during each session. Data were analyzed using a one-sample Student’s *t*-test comparing the mean exploration percentage time for each object with the chance value of 50%. Animals that recognize the familiar object as such (i.e., learn the task) explore the novel object >50% of the total time.

*Morris Water Maze (MWM):* The apparatus used for the water maze task was a circular tank (1.2 m diameter) filled with water maintained at 20 ± 0.5 °C.  The tank was located in a test room containing prominent visual cues.  Mice were trained to swim to a 11 cm diameter circular platform submerged 1.5 cm beneath the surface of the water and invisible to the mice while swimming.  The platform was located in a fixed position, equidistant from the center and the wall of the tank.  Mice were subjected to four training trials per day (inter-trial interval, 10 min).  On each trial, mice were placed into the tank at one of four designated start points in a pseudorandom order.  Mice were allowed to find and escape onto the submerged platform.  If they failed to find the platform within 60 sec, they were manually guided to the platform and allowed to remain for 10 sec.  Mice were trained for four consecutive days.  The probe trial was assessed 24 hours after the last training session and consisted of a 60 sec free swim in the pool without the platform.  Data were collected using the ANY-maze behavioral tracking software (Stoelting).

*Rotarod:* The test was performed in a mouse rotarod apparatus (Insight Ltda.,Brazil), as previously described. Briefly, mice were individually placed in the apparatus floor for 3 minutes followed by a 2-min habituation session to the cylinder rod. The test phase consisted of tree trials (inter-trial interval, 60 min) in which animals were placed on the top of the rod rotating at increasing speed (minimal speed 16 rpm, maximal speed 36 rpm with acceleration rate 3.7 rpm). Latency to fall was recorded for a 5 min period, and results are expressed as average latency in the test phase.

**Tissue collection**

Animals were anesthetized (90 mg/kg ketamine and 4.5 mg/kg xylazine, i.p.) before perfusion with ice-cold PBS at early and late phases of the model. Hippocampal tissues were dissected immediately after perfusion, frozen in liquid nitrogen and stored at -80ºC before RNA extraction. For immunofluorescence studies, perfusion was performed with 4% PFA, and brains were fixed for 24 h before paraffin processing.

**Cell culture and treatments**

Primary neuronal cortical culture was prepared as previously described in Diniz 2012. Briefly, dissociated cerebral cortices were harvested from embryonic day 14 Swiss mice and cultured in neurobasal medium (Invitrogen) supplemented with B-27, penicillin, streptomycin, l-glutamine, fungizone and cytosine arabinose, and maintained at 37ºC with 5% CO_2_. Neurons were seeded at a density of 50.000-150.000 neurons/well on a 13mm diameter poly-D-lysine-coated well (10µg/mL; Sigma). One week after dissociation, neuronal cell cultures were treated with PBS or spike protein (1µg/mL) for 24 h. Later, cells were fixed in 4% PFA, 6% sucrose in PBS for 10 min before immunocytochemistry assay.

The murine BV-2 cell line was cultured in DMEM supplemented with 10%  FBS, and 1% streptomycin/penicillin, and seeded at a density of 100.000 cells/well on a 13mm diameter poly-D-lysine-coated well. Next, cells were treated with PBS or spike protein (1µg/mL) for 24 h and fixed as mentioned above.

**RNA extraction and qPCR**

RNA extraction of hippocampal tissue and cell cultures was performed using Trizol® reagent (Invitrogen), in accordance with manufacturer’s instructions. Sample concentration and purity was assessed using a NanoDrop 1000 spectrophotometer. (ThermoScientific). Only preparations with absorbance ratios >1.8 and no signs of RNA degradation were used. One μg of total RNA was reverse transcribed using the High-Capacity cDNA Reverse Transcription Kit (Applied Biosystems), according to the manufacturer's instructions. qPCR was performed using a QuantStudio 5 PCR system (Applied Biosystems) with reactions performed in triplicate. Briefly, qPCRs were run using Power SYBR Green PCR Master Mix (Life Technologies), and 10 ng of template cDNA in a 10 μL reaction volume. The primers used are listed in Supplementary Table 2. Cycle threshold (Ct) values were normalized to a control gene (β-actin) and analyzed using the ΔΔCt method to generate fold change values (2^–ΔΔCT^).

**Immunofluorescence assay**

Slides containing the hippocampal formation were deparaffinized, and antigen retrieval was carried out by incubation in citrate buffer solution (pH 6.0) at 95ºC for 40 min. Afterwards, permeabilization was performed with 0.025% Triton in PBS, followed by incubation with blocking buffer (PBS containing 0.025% Triton, 3% BSA, and 5% normal goat serum) for 2 h. Next, slides were incubated overnight with primary antibodies against IBA-1 (WAKO; 1:800#019-19741), synaptophysin (Vector Laboratories; 1:200 #S285), Homer-1 (Abcam; 1:100 #184955), or GFAP (Sigma; 1:500 #G3893). For immunocytochemistry, wells were washed three times with PBS, and incubated for 1 h with blocking buffer, followed by overnight incubation with primary antibodies against β3-tubulin (Promega; 1:1000 #G712A), Iba-1 (1:1000), synaptophysin or Homer-1. For visualization, sections or wells were incubated with AlexaFluor 488- or 546-conjugated secondary antibodies for 2 h at room temperature, washed with PBS and mounted in Fluoroshield with DAPI (Sigma).

The β3-tubulin immunoreactivity in cortical neurons, Iba-1 immunoreactivity in BV-2 cells, as well as microglia density and morphology in Iba-1 immunostained brain sections were photographed using a Slight DS-5-M1 digital camera (Nikon,Melville,NY) connected to an epifluorescence Nikon Eclipse 50i light microscope, under a 20 or 40x objective. Microglia morphology was assessed evaluating the number of branches emanating from their soma (Lopez-Rodrigues et al., 2015). Briefly, type I and type II cells were described as surveillant microglia and present smaller soma and less than 5 thin branches. Type III, IV and V microglia are characterized as reactive microglia, and present more than 4 branches, and thicker branches and bigger soma are observed (Lopez-Rodrigues et al., 2015). Optical density for β3-tubulin and Iba-1 was measured using ImageJ v1.53 and normalized by total DAPI stains. Pyknotic nuclei were analyzed using DAPI stains with 400x magnification and normalized by the total DAPI-stained nuclei observed. In order to determine the synaptic density or synapse engulfment, 9-12 confocal z-stacks (0,35um/z-stack) were obtained using a Leica TSE-SPE3 confocal microscope (Figure 1) or using a Zeiss Cell Observer Spining Disk Confocal microscope (Figure 4) at 630x magnification, and each z-stack was individually analyzed using the ImageJ v1.53 plugin SynQuant automated synapse counter (Wang et al., 2020). Quantitative colocalization of post- (Homer-1) and presynaptic (synaptophysin) markers, or Iba-1 and synaptophysin in control mice were used to normalize the ratio of preserved synaptic puncta and synaptic engulfment, respectively.

**Enzyme-linked immunosorbent assay (ELISA)**

For cytokine measurements, hippocampus was homogenized in cold RIPA buffer (150 mM NaCl, 1% Triton X-100, 0.5% sodium deoxycholate, 0.1% SDS, 50 mM Tris Base, 2 mM PMSF, pH 8), and supernatant was collected after centrifugation at 14,000 *g* for 10 min at 4°C.  Protein concentration as determined using the [BCA Protein Assay](https://www.thermofisher.com/order/catalog/product/23227) (Thermo Scientific). Samples diluted 1:10 in RIPA buffer were used for the detection of TNF (BD Biosciences) and IL1β (R&D Systems) by ELISA according to manufacturer's instructions. Results were expressed as pg/µg protein.

**Neurofilament light chain (NFL) measurements**

Mouse plasma NFL concentration was measured in triplicate using ultra-sensitive single molecule array (Simoa) technique on the Simoa SR-X™ Analyzer, using Simoa NF-Light Advantage according to the manufacturer's instructions (Quanterix). Briefly, plasma samples were thawed at room temperature for one hour and then centrifuged at 10,000 RCF for 5 minutes at 24°C. Samples were diluted 1:4 with sample diluent and applied to the plate in duplicate. Paramagnetic beads coated with capture anti-NFL were incubated with a biotinylated anti-NfL detection antibody, followed by incubation with a streptavidin-β-galactosidase complex. A fluorescent signal proportional to the concentration of NfL was generated after the addition of the substrate resorufin β-D-galactopyranoside. Controls were used to validate the detection limit of 0.0552 pg/mL. All coefficients of variance (CVs) of duplicate measurements were below 20%.

**Study population and cognitive assessment**

Post-COVID-19 outpatients were evaluated between December 2020 and July 2021 by a multidisciplinary team of neurologists and neuropsychologists at the Gaffrée and Guinle University Hospital (Rio de Janeiro, Brazil). Inclusion criteria included: COVID-19 diagnosis confirmed by PCR or serological diagnosis, fulfilling criteria of mild disease (not require hospitalization and symptoms that not included dyspnea); assessment performed at least 15 days after the end of symptoms, blood collection and neurocognitive evaluation consent. Exclusion criteria included: age under 18 years old; individuals with previously known cognitive impairment or other neuropsychiatrist disorders that could interfere with the test results. All study subjects had their detailed clinical history recorded and were subjected to complete physical and neurological examination. This work was approved by the Brazilian Ethics Committee (CONEP, CAAE 33659620.1.1001.5258). All participants signed the informed consent term, agreeing to participate in this research.

Neurocognitive status was assessed using the Symbol Digit Modalities Test (SDMT), a screening test developed to identify individuals with cognitive impairment through the domains of attention, processing speed and motor skills. The raw score of the SDMT is converted to scaled scores (M = 10, SD = 3) using the cumulative frequency distribution of the test in order to normalize test score distributions. The resulting scaled scores is regressed on age, age-squared, sex, and education. After the evaluation, the participants of the study were divided into two main subgroups, “with cognitive deficit” and “without cognitive deficit”.

**Sample collection and genotyping**

Blood samples were collected and centrifuged at 1.500 *g* at 4 °C for 15 min to separate the buffy coat from plasma. Genomic DNA (gDNA) was extracted using the PureLink Genomic DNA Mini Kit (ThermoFisher Scientific). The quality of the gDNA was determined using NanoDrop 2000 (ThermoFisher Scientific) followed by quantification using the Qubit dsDNA HS Assay Kit (ThermoFisher Scientific) and Qubit Fluorometer 3.0 (Thermo Fisher Scientific).

The *TLR4* -2604G>A (rs10759931) and *TLR4* - 2272A>G (rs2737190) variants were genotyped with allelic discrimination using TaqMan qPCR system (ThermoFisher Scientific). The probes were produced by Applied Biosystems [rs10759931 (C___2704046_10) and rs2737190 (C___2704047_10)]. Briefly, genotyping was performed in a 20 µL reaction mixture containing 10 ng DNA, TaqMan Universal PCR Master Mix (1X), Probe TaqMan Gene Expression Assay (1X), and DNAse-free water for the final volume. The reaction was carried out in the following conditions: an UNG incubation step of 2 min at 50 ^◦^C, polymerase activation for 10 min at 95 ^◦^C, followed by 40 cycles of 15 s at 95 ^◦^C for denaturation and 60 s at 60 ^◦^C for annealing/extension. The amplification and reading of the plates were performed in the QuantStudio 5 Real-Time PCR System (Applied Biosystems).

**Illustrations**

Illustrations in figures 1A, 3G and 4C and M were created using  *MindtheGraph* (www.mindthegraph.com; under FFD subscription) and subsequently modified (free culture Creative Commons license).

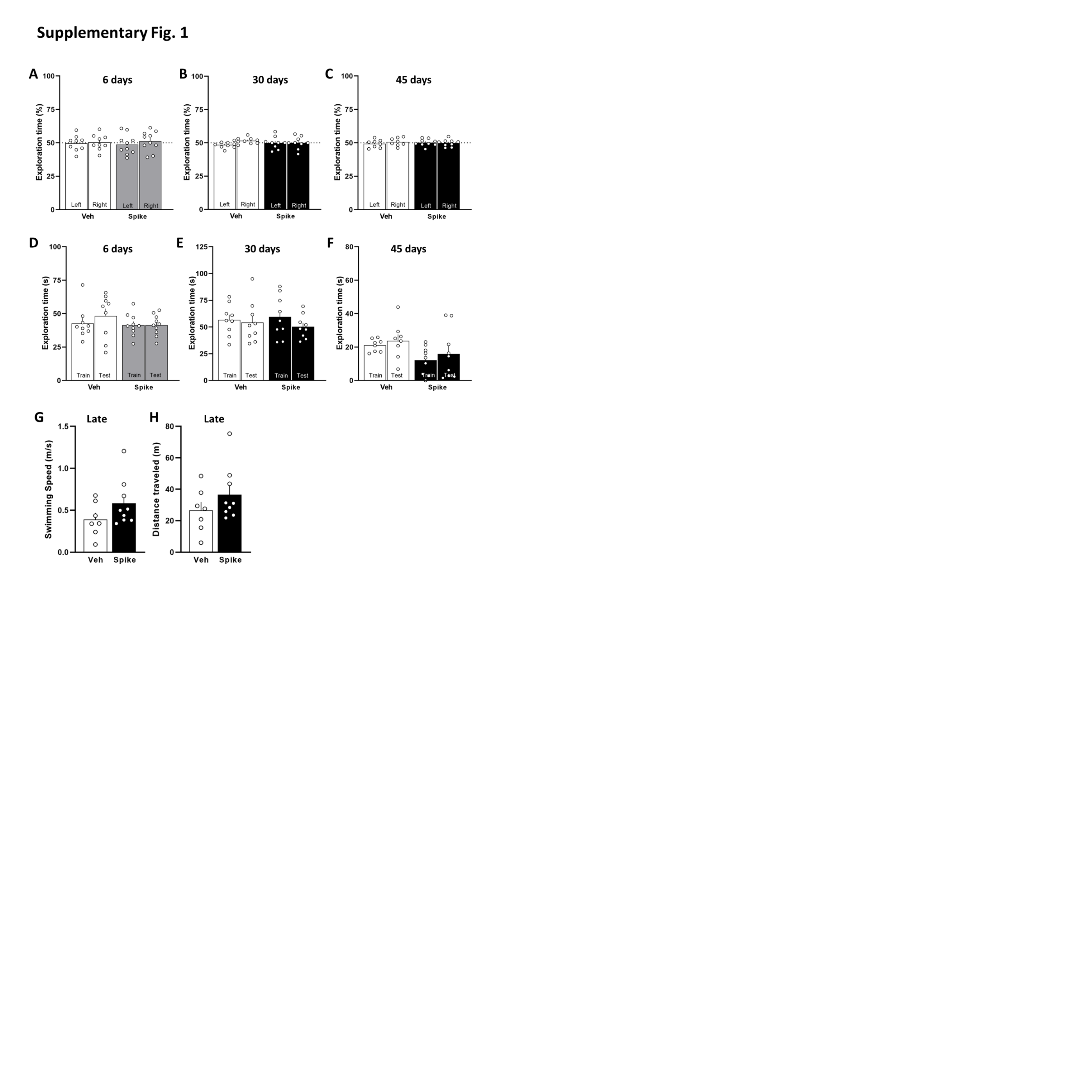

**Fig. S1 Controls of behavioral tests and serological analyses.** Intracerebroventricular (icv) infusion of Spike protein had no effect on innate preferences for the objects during the training session (**A to C**), or exploratory activity (**D to F**) during the test session of NOR at 6, 30 and 45 days after protein infusion (**A to F**; *N* = 8-10 mice per group). Spike protein does not change swimming speed (**G**) or total distance traveled (**H**) in the Morris Water Maze (*N* = 7-9 mice per group). Bars represent means ± SEM.

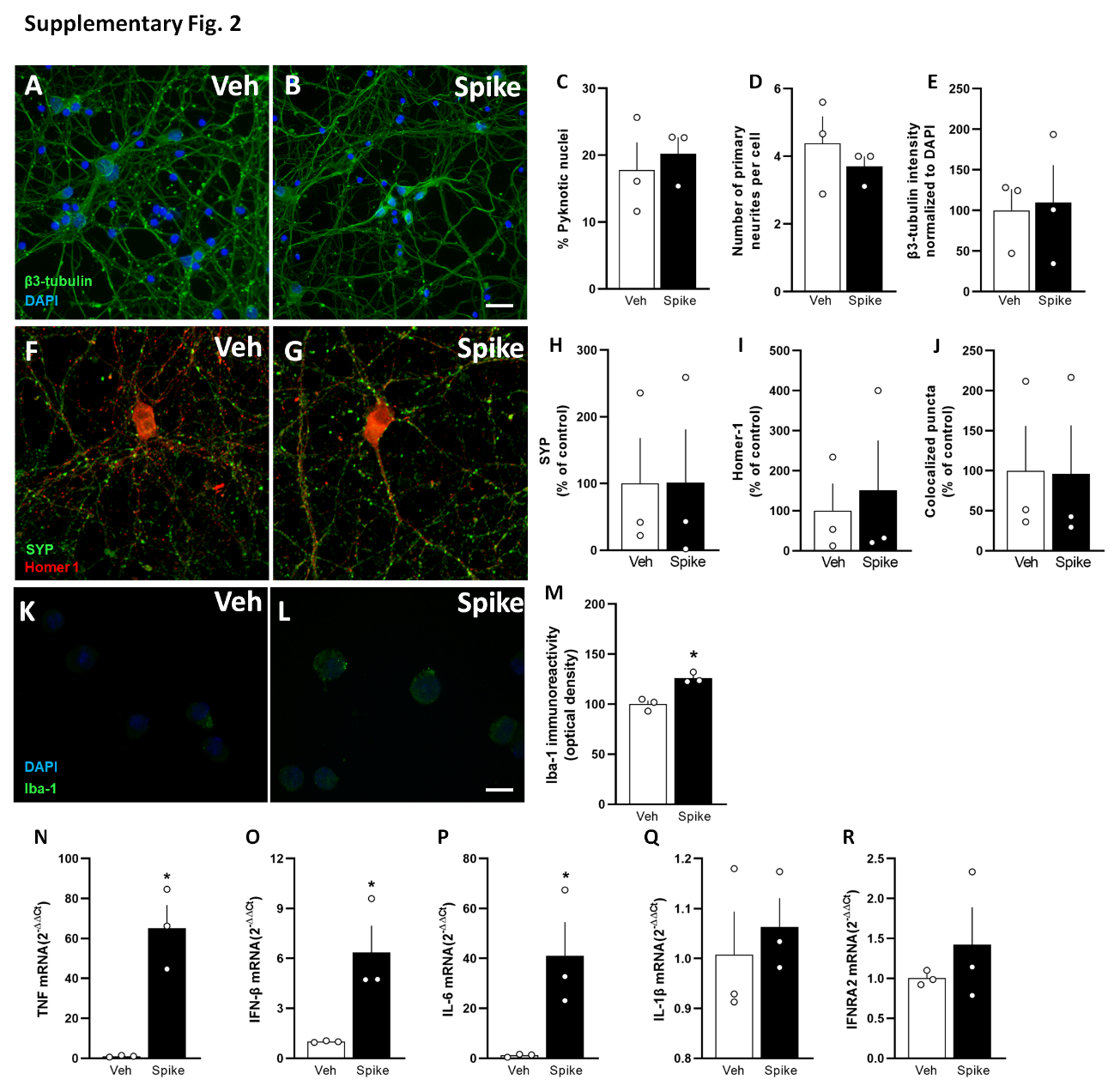

**Supplementary Fig. 2 Microglial and neuronal culture**. Cultured primary cortical neurons were incubated with SARS-CoV-2 spike protein (Spike) or vehicle (Veh) during 24h, and analyzed by immunocytochemistry. (**a-b, f-g**). Spike protein does not induce changes in the number of pyknotic nuclei (**c**), primary dendrites (**d**) and β3-tubulin intensity (**e**). (**f-j**) Spike protein also induces no difference in the number of synapses in cortical neurons, as demonstrated by double immunostaining for Homer-1 (**h**) and synaptophysin (SYP) (**i**) followed by determination of the colocalization rate (**j**). (**a-j**; *N* = 5 experiments with independent neuron cultures). Representative images of IBA-1 immunoreactivity in BV-2 cells incubated with Veh (**k**) or Spike protein (**l**). bar = 20µm. **m-r,** BV2 cells were treated with SARS-CoV-2 spike protein (Spike) or vehicle (Veh) during 24h, and analyzed by immunocytochemistry or qPCR. Iba-1 immunoreactivity (**m** t=5.567, **p* = 0.0051), mRNA levels of TNF (**n** t=5.557, **p* = 0.0051), IFN-β (**o** t=3.307, **p* = 0.0297), IL-6 (**p** t=2.968, **p* = 0.0412), IL-1β (**q**), and IFNAR2 (**r**); Student’s *t*-test; *N* = 3. Bars represent means ± SEM.

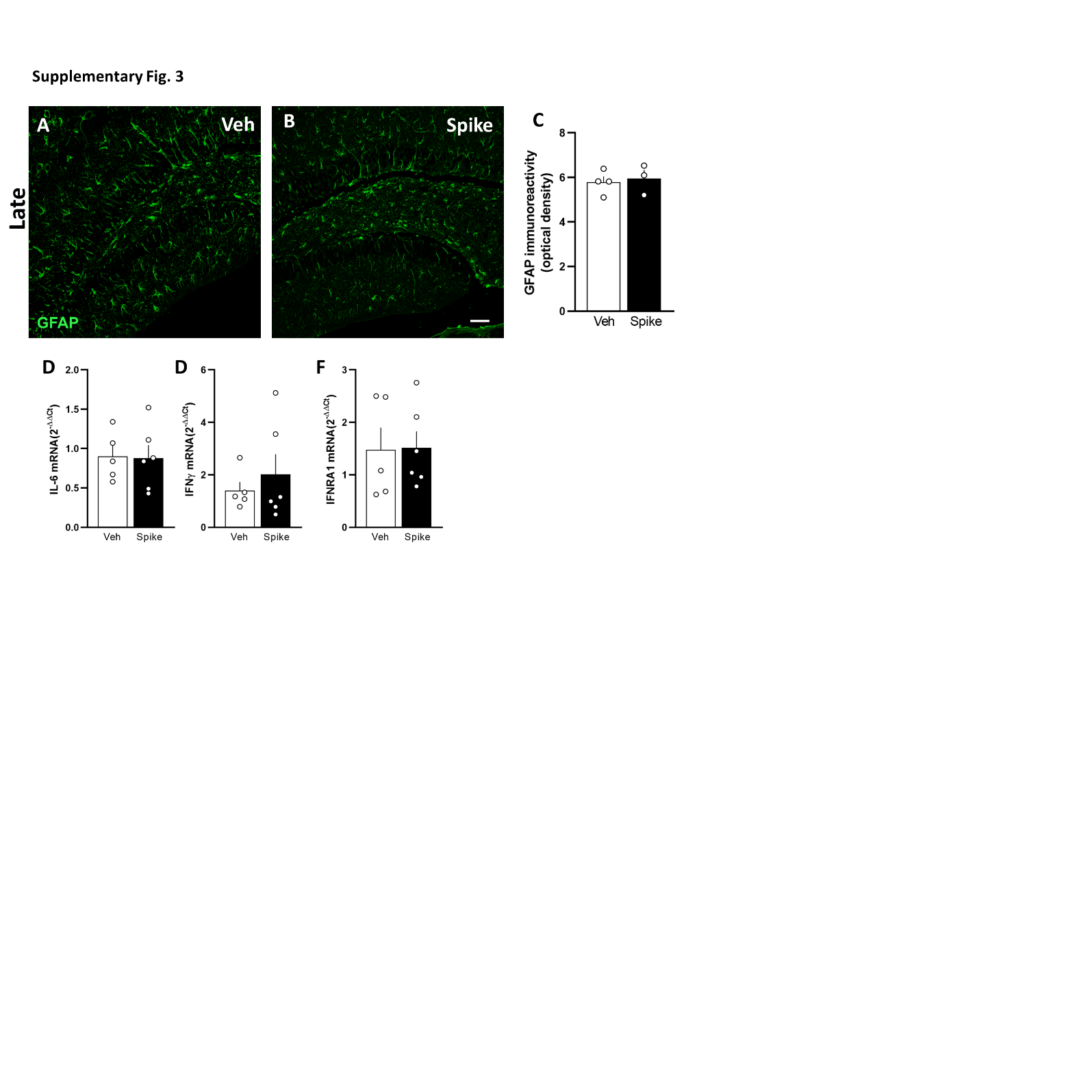

**Fig. S3** **GFAP immunohistochemistry and qPCR analysis.** (**A to C**) Spike protein had no effect on GFAP immunoreactivity in the hippocampus of mice, when evaluated later after protein brain infusion (**A to C**; *N* = 3-4 mice per group)), nor change the mRNA levels of IL-6 (**D**), IFNγ (**E**) and IFNAR1 (**F**) (*N* = 5-6 mice per group). Scale bar = 50 µm. Bars represent means ± SEM.

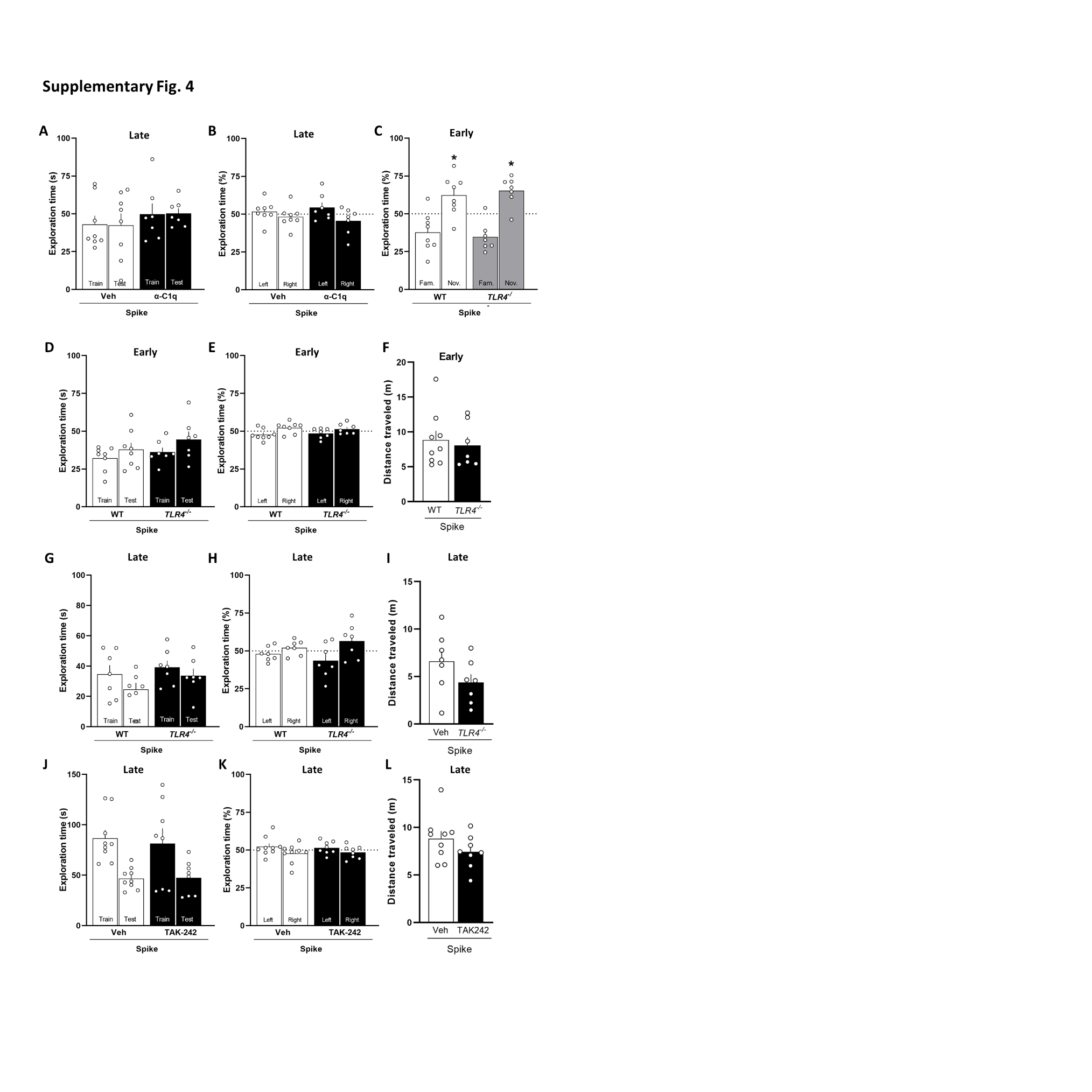

**Fig. S4** **Controls of behavioral tests.** Intracerebroventricular (icv) infusion of Spike protein had no effect on exploratory activity during the test session (**A, D, G** and **J**) or innate preferences for the objects during the training session (**B, E, H** and **K**) in the NOR (*N* = 7-9 mice per group). (**C**) Spike protein does not impair object recognition memory in WT and TLR4^-/-^ mice, early after protein infusion (t=2.66 *p=0.0323 for WT and t=4.18 *p=0.0058 for TLR4^-/-^); one-sample Student’s *t*-test compared to the chance level of 50% (*N* = 7-8 mice per group). Genetic absence (**F** and **I**) or blockade of TLR4 (**L**) does not affect total distance traveled in the open field arena (*N* = 7-9 mice per group). Bars represent means ± SEM.

**Supplementary table 1. Participant demographics of the study sample.**

| **Sample demographics** | **Number of individuals (%)**  **(total *n* = 86)** |
| --- | --- |
| Sex |  |
| Female | 70 (81.4%) |
| Male | 16 (18.6%) |
| Age (years)^a^ | 45.6 (19-71) |
| Time between onset of clinical symptoms and cognitive assessment (months)^a^ | 6.85 (1-15) |
| Education (years)^a^ | 17.02 (5-28) |
| Comorbidities |  |
| 1. None | 40 (45.5%) |
| 2. Obesity | 19 (22.1%) |
| 3. Hypertension | 17 (19.7%) |
| 4. Diabetes | 10 (11.6%) |

a= mean (range)

| **Target gene** | **Forward primer** | **Reverse primer** |
| --- | --- | --- |
| β-Actin | GCCCTGAGGCTCTTTTCCAG | TGCCACAGGATTCCATACCC |
| TNF | CCCTCACACTCAGATCATCTTCT | GCTACGACGACGTGGGCTACAG |
| IFNβ | CACAGCCCTCTCCATCAACTA | CATTTCCGAATGTTCGTCCT |
| Il6 | GCTACCAAACTGGATATAATCAGGA | CCAGGTAGCTATGGTACTCCAGAA |
| IL1-β | GTAATGAAAGACGGCACACC- | ATTAGAAACAGTCCAGCCCA- |
| IFNAR1 | CTGGTCTGTGAGCTGTACTT | TCCCCGCAGTATTGATGAGT |
| IFNAR2 | CTATCGTAATGCTGAAACGG | CGTAATTCCACAGTCTCTTCT |
| IFNγ | AGCAACAGCAAGGCGAAAA | CTGGACCTGTGGGTTGTTGA |
| C1q | CTCAGGGATGGCTGGTGGCC | CCTTTGAGACCCGGCCTCCCC |
| TLR4 | GTCAGTGTGATTGTGGTATCC | ACCCAGTCCTCATTCTGACTC |

**Supplementary Table 2:** List of primers used in qPCR analyses.
